# Proteomic mapping of novel tubulin post-translational modifications in *Trypanosoma cruzi* cytoskeleton

**DOI:** 10.64898/2026.04.09.717369

**Authors:** Gonzalo Martinez Peralta, Daiana Baldelomar, Lucila Baldasseroni, Azul de Hernandez, Esteban Serra, Victoria Lucía Alonso

## Abstract

Microtubules play central roles in the organization and morphology of trypanosomatid parasites, forming highly specialized cytoskeletal structures such as the subpellicular corset, the flagellar axoneme, and the mitotic spindle. Functional specialization of MTs is proposed to be regulated by the “tubulin code”, which is defined by the combination of different α- and β-tubulin isotypes, a set of post-translational modifications and specific MT-binding proteins. Although multiple tubulin post-translational modifications have been described in trypanosomatids using specific antibodies or mass spectrometry, to date no comprehensive mapping has been reported in *Trypanosoma cruzi*, the causative agent of Chagas Disease. In the present work, we performed a high-resolution proteomic analysis of PTMs present in α- and β-tubulin subunits of the *T. cruzi* Dm28c strain, using tubulin-enriched extracts obtained by *in vitro* polymerization. Multiple high-confidence PTMs were identified by LC–MS/MS, including acetylation, methylation, phosphorylation, and polyglutamylation, for which many modified amino acids had not been previously reported in trypanosomatids. Structural mapping of these modifications onto a predicted α/β-tubulin heterodimer showed that most modified residues are located in solvent-exposed regions of the protein. Together, these findings provide the first high-confidence proteomic landscape of tubulin PTMs in *T. cruzi* and support the existence of a complex tubulin code contributing to microtubule regulation in this parasite.

## INTRODUCTION

The cytoskeleton of trypanosomatids is formed by an extensive network of microtubules (MTs) that plays a central role in parasite morphology, polarity, cellular organization, cell division and differentiation during its life cycle. In *Trypanosoma cruzi* (and Trypanosomatids), the cell body is supported by a highly ordered array of subpellicular MTs located below the plasma membrane. This structure, which resembles a corset, confers mechanical rigidity and defines the characteristic morphology of the parasite. The association between subpellicular MTs and the plasma membrane is remarkably stable and remains intact even after cell lysis and isolation of membrane fractions (Vidal & Souza, 2017). Electron microscopy studies have shown that subpellicular MTs are arranged in parallel arrays with regular spacing between individual MTs and relative to the plasma membrane. MT-free regions are only present at the flagellar pocket and the cytostome-cytopharynx complex in *T. cruzi*, allowing essential endocytic and exocytic processes to occur in these specialized domains (De Souza, 2009).

In addition to the subpellicular MT corset, trypanosomatids possess several specialized cytoskeletal elements. The parasite flagellum is essential for viability and contributes to motility as well as processes related to morphogenesis, cell division, transmission, and host interaction (Langousis & Hill, 2014; Zuma et al., 2021). In this structure, the flagellar axoneme follows a conserved 9 + 2 arrangement typical of eukaryotic flagella and drives the undulatory movements that propel the parasite. The mitotic spindle is composed of highly dynamic MTs capable of rapid assembly and disassembly during mitosis (Gull, 1999). The remarkable structural organization of the trypanosomatid cytoskeleton and the presence of single-copy organelles make these parasites valuable model systems for studying cytoskeletal organization and microtubule biology.

MT properties are influenced by the diversity of tubulin isotypes encoded by different α- and β-tubulin genes and by a wide range of post-translational modifications (PTMs). These modifications include acetylation, tyrosination/detyrosination, phosphorylation, glutamylation, and glycylation, and can occur both within the globular domains and within the flexible C-terminal tails of tubulins. The combination of tubulin isotypes and PTMs generates multiple tubulin isoforms that contribute to the functional specialization of MTs. This regulatory mechanism is referred to as the “tubulin code” (Gadadhar et al., 2017; Janke & Magiera, 2020).

In *T. cruzi*, the regulatory mechanisms that involve tubulin PTMs are starting to be explored. Acetylation of α-tubulin at lysine 40 (K40) is one of the most characterized PTMs and has been associated with microtubule stability in different organisms (Li & Yang, 2015; Lu, 2025; Perdiz et al., 2011). In *T. cruzi*, the α-tubulin acetyltransferase responsible for this modification (*Tc*ATAT) has been identified and shown to play a key role in parasite growth and cell cycle progression (Alonso et al., 2021).

Other enzymes potentially involved in regulating tubulin PTMs have been described in trypanosomatids. Homologues of tubulin tyrosine ligase-like enzymes (TTLLs), responsible for polyglutamylation and polyglycylation in other eukaryotes, are present in kinetoplastid genomes, together with cytosolic carboxypeptidases that can remove glutamate side chains (Casanova et al., 2015; Jentzsch et al., 2020). Furthermore, deacetylase enzymes belonging to the HDAC and Sirtuin families have been implicated in regulating tubulin acetylation levels (De Oliveira Santos et al., 2019; de Oliveira Santos et al., 2021; Ritagliati et al., 2015). These observations suggest that a complex enzymatic machinery capable of establishing and reversing tubulin PTMs may operate in trypanosomatids.

Despite these advances, a comprehensive characterization of the tubulin PTM landscape in *T. cruzi* (and other trypanosomatids) is still lacking. Most studies have focused on individual PTMs or relied on antibody-based detection approaches, leaving the global repertoire of tubulin PTMs largely unexplored. Moreover, the sequence divergence of trypanosomatid tubulins, particularly within the C-terminal tails, complicates the prediction of PTMs based solely on homology with other eukaryotic systems.

In the present study, we performed a proteomic analysis to identify PTMs present in α- and β-tubulin from the *T. cruzi* Dm28c strain. Using tubulin-enriched extracts obtained through *in vitro* polymerization and high-resolution mass spectrometry, we identified multiple modification sites distributed across both tubulin subunits. These results provide the first systematic mapping of tubulin PTMs in *T. cruzi* and establish a valuable framework for future studies aimed at understanding the regulation and functional specialization of parasite MTs.

## MATERIALS AND METHODS

### T. cruzi epimastigotes culture conditions

The *T. cruzi* Dm28c strain was used in this study, which belongs to the TcI group (Zingales et al., 2012). Epimastigotes were cultured at 28 °C in Liver Infusion Tryptose (LIT) medium, composed of 5 g/L liver infusion, 5 g/L tryptose, 4 g/L NaCl, 0.4 g/L KCl, 8 g/L Na_2_HPO_4_, 4 g/L glucose, and supplemented with 5 µM hemin and 10% (v/v) fetal bovine serum (FBS). The parasites were kept in an exponential growth phase by subculturing every 3 days.

### Purification of *in vitro* polymerized tubulin from epimastigotes

The following protocol was adapted from MacRae and Gull (Macrae & Gull, 1990). Briefly, 200 mL of parasites in exponential phase (30 × 10^6^ parasites/mL) were harvested by centrifugation at 2500 g for 10 min at 16 °C. The resulting pellet was washed twice with 10 mL of PEM buffer (100 mM PIPES, 1 mM EGTA, 1 mM MgSO_4_, pH 6.9). The pellet was resuspended in 5 mL of PEM buffer supplemented with protease inhibitors (1 mM AEBSF, 2 µg/mL aprotinin, 1 µM bestatin, 10 µM E-64, 10 µM leupeptin, 1 µM pepstatin A, and 1 mM PMSF), hereafter referred to as PEM + inhibitors. Cells were sonicated on ice with six pulses of 20 s at 30% amplitude. The resulting lysate, referred to as the total extract (TE), was incubated on ice for 30 min and subsequently centrifuged in an ultracentrifuge (Beckman Coulter Optima XL-100K, Ti90 rotor) at 1 × 10^5^ g for 1h at 4 °C. The supernatant containing soluble MTs (S1) and the pellet lacking MTs (P0) were collected separately. The P0 fraction was resuspended in 1 mL PEM + inhibitors and stored at −80 °C. To the S1 fraction, 10 µL of Taxol (1 mM) and 250 µL of GTP (10 mg/mL) were added to induce tubulin polymerization. The mixture was incubated at 30 °C for 1h with gentle agitation. Samples were then centrifuged at 1 × 10^5^ g for 30 min at 25 °C. The supernatant lacking MTs (S2) and the pellet containing polymerized MTs (P-MT) were collected separately. The P-MT pellet was resuspended in 300 µL of PEM + inhibitors and stored at −80 °C. Aliquots (40 µL) of each fraction were collected at each step for SDS-PAGE and Western blot analysis.

### SDS-Page and western blot

Protein extracts were separated by SDS-PAGE on 12% polyacrylamide gels under denaturing conditions. Samples were mixed with Laemmli loading buffer, boiled for 5 min, and centrifuged to remove insoluble material prior to loading. Electrophoresis was performed in Tris-Glycine-SDS running buffer and gels were stained with Coomassie Blue G-250.

For Western blot analysis, proteins were transferred onto nitrocellulose membranes using a Mini Trans-Blot system (Bio-Rad) according to the manufacturer’s instructions. Transfer efficiency was verified by Ponceau S staining. Membranes were blocked with 5% non-fat milk in PBS and incubated overnight at 4 °C with primary antibodies diluted in PBS containing 0.5% Tween 20 (PBS-T). The following antibodies were used: mouse monoclonal anti-α-tubulin (1:2000, clone DM1A, Cell Signalling Technology Cat# 3873, RRID:AB_1904178), mouse monoclonal anti-acetylated α-tubulin (1:2000, clone 6-11B-1, Sigma-Aldrich Cat# MABT868, RRID:AB_2819178), mouse monoclonal anti-β-tubulin (1:2000, clone KMX-1, Millipore Cat# MAB3408, RRID:AB_94650), rat monoclonal anti-tyrosinated α-tubulin (1:2000, clone YL1/2, Millipore Cat# MAB1864-I, RRID:AB_2890657), and rabbit polyclonal anti-methylated Lysine (1:1000, Abcam, Cat#ab23366, RRI D: AB_447401). After washing, membranes were incubated with HRP-conjugated secondary antibodies: goat anti-rat IgG (1:20000, Sigma-Aldrich Cat# AP136P, RRID:AB_11214444) and rabbit anti-mouse IgG (1:20000, Sigma-Aldrich Cat# 12-349, RRID:AB_390192) and immunoreactive bands were detected using the ECL Plus system (GE Healthcare). Signals were visualized using an GE Amersham Imager 600 (RRID:SCR_021853).

### Indirect immunofluorescence

Exponentially growing epimastigotes were centrifuged, washed twice in PBS, settled on polylisine-coated coverslips and fixed with 4% para-formaldehyde in PBS at room temperature for 20 minutes. Fixed parasites were washed with PBS and permeabilized with 0.1% Triton X-100 in PBS for 10 minutes. After washing with PBS, parasites were incubated with the appropriate primary antibody diluted in 5% BSA in PBS for 2 hours at room temperature. Primary antibodies used were: rabbit polyclonal anti-Methylated Lysine (1:100, Abcam, Cat#ab23366, RRID: AB_447401) and mouse monoclonal anti-acetylated α-tubulin (1:200, clone 6-11B-1, Sigma-Aldrich Cat# MABT868, RRID:AB_2819178). In colocalization experiments both antibodies were incubated together. Non-bound antibodies were washed with 0.01% Tween 20 in PBS and then the slides were incubated with fluorescent-conjugated goat anti-mouse Alexa-555 (1:500, Thermo Fisher Scientific Cat# A-21422, RRID:AB_2535844) and goat anti-rabbit FITC IgG antibodies (1:500, Sigma-Aldrich Cat# F1262, RRID:AB_259430) and 2 µg/mL of DAPI for 1 hour. The slides were washed with 0.01% Tween 20 in PBS and finally mounted with VectaShield (Vector Laboratories). Images were acquired with a confocal Zeiss LSM880. ImageJ software were used to process all images. Colocalization analysis was performed using the Coloc2 plugin implemented in Fiji/ImageJ. Pearson’s correlation coefficient and Costes’ randomization test were calculated using the default parameters using three images with approximately 20 parasites each.

### Sample preparation for LC-MS

An SDS-PAGE gel was run loading 20 µg of the P-MT sample and stained with colloidal Coomassie blue. The bands corresponding to α-tubulin and β-tubulin were excised and cut into several fragments, which were processed following the protocol for in-gel tryptic digestion (Link & LaBaer, 2009). This process was performed at the Mass Spectrometry Unit of the Institute of Molecular and Cellular Biology of Rosario (UEM-IBR), where LC-MS analysis was subsequently carried out.

### LC-MS analysis

Peptide separations were carried out on a Thermo Scientific Ultimate 3000 RSLCNano UPLC system (RRID:SCR_026145) using a C18 nano column (15 cm, Thermo Fisher Scientific). The mobile phase flow rate was 300 nL/min using 0.1% formic acid in water (solvent A) and 0.1% formic acid and 100% acetonitrile (solvent B). The gradient profile was set as follows: 4-30% solvent B for 64 min, 30%-80% solvent B for 7 min and 80% solvent B for 1 min. 3 microliters of the sample were injected. MS analysis was performed using a Thermo Scientific Q Exactive HF Orbitrap LC-MS/MS System (RRID:SCR_020545). For ionization, 1.9 kV of liquid junction voltage and 300 °C capillary temperature was used. The full scan method employed a m/z 200–2500 mass selection, an Orbitrap resolution of 120000 (at m/z 200), a target automatic gain control (AGC) value of 1e6, and maximum injection times of 100 ms. After the survey scan, the 15 most intense precursor ions were selected for MS/MS fragmentation. Fragmentation was performed with a normalized collision energy of 27 eV and MS/MS scans were acquired with a dynamic first mass, AGC target was 5e5, resolution of 30000 (at m/z 200), isolation window of 1.4 m/z units and maximum IT was 55 ms. Charge state screening was enabled to reject unassigned, singly charged, and equal or more than six protonated ions. A dynamic exclusion time of 25 s was used to discriminate against previously selected ions.

### MS data analysis

For the analysis of tubulin PTM data, two different software platforms were used: Proteome Discoverer (v2.4.1.15) (Thermo Fisher Scientific, RRID:SCR_014477) and MaxQuant (v2.7.5.0) (Max Planck Institute of Biochemistry, Germany, RRID:SCR_014485), the latter being open-source.

MS data were analysed with Proteome Discoverer (version 2.4.1.15) using standardized workflows (Orsburn, 2021). Mass spectra *.raw files were searched against a database from *T. cruzi Dm28c* (UP000017861), supplemented with the specific sequences for α-tubulin (C4B63_51g126) and β-tubulin (C4B63_22g172) from *T. cruzi* Dm28c 2018 strain obtained from TritrypDB (www.tritripdb.org)(Berná et al., 2018). Precursor and fragment mass tolerance were set to 10 ppm and 0.02 Da, respectively, allowing 2 missed cleavages.

The following dynamic PTMs were included in the search parameters: oxidation (+15.995 Da; M), acetylation (+42.011 Da; K, R), methylation (+14.016 Da; K), phosphorylation (+79.966 Da; S, T, Y), and tyrosination (+163.063 Da; E). In addition, dynamic PTMs at the N-terminal end were considered, including N-terminal acetylation (+42.011 Da), methionine loss (−131.040 Da; M), and the combined modification methionine loss + acetylation (−89.030 Da; M). Carbamidomethylation (+57.021 Da; C) was set as a static modification due to the iodoacetamide treatment.

To specifically explore the presence of polyglutamylation and polyglycylation in the C-terminal tails of α- and β-tubulin, the raw data were also reprocessed using MaxQuant (v2.7.5.0) with the Andromeda search engine. For this targeted analysis, a reduced .fasta database containing only the α- and β-tubulin sequences of *T. cruzi* Dm28c was used. PTMs were manually added to the modifications.local.xml file in MaxQuant.

For polyglutamylation, the following PTMs were defined: polyGlu_1 (C(5)H(7)NO(3), +129.0426 Da; E), polyGlu_2 (C(10)H(14)N(2)O(6), +258.0852 Da; E), and polyGlu_3 (C(15)H(21)N(3)O(9), +387.1278 Da; E). For polyglycylation, the following PTMs were defined: polyGly_1 (C(2)H(3)NO, +57.0215 Da; E), polyGly_2 (C(4)H(6)N(2)O(2), +114.0429 Da; E), and polyGly_3 (C(6)H(9)N(3)O(3), +171.0644 Da; E). In all cases, the modifications were restricted to glutamate residues (E) and defined as standards.

Polyglutamylation and polyglycylation analyses were performed in separate runs in order to reduce computational complexity. In both cases, carbamidomethylation was selected as a fixed modification, while methionine oxidation, N-terminal acetylation, and the different states of polyglutamylation (or polyglycylation) were defined as variable PTMs. The maximum number of variable modifications per peptide was set to two, and label-free quantification (LFQ) was not performed.

Trypsin was selected as the digestion enzyme, allowing up to two missed cleavage sites per peptide. Finally, the evidence.txt output file was used to filter high-confidence peptides corresponding to the C-terminal tails. Only peptides meeting the following criteria were retained: posterior error probability (PEP) ≤ 0.01, mass error between −2 and 2 ppm, and at least two MS/MS spectra (MS/MS count ≥ 2). Only high-confidence PTMs were considered for further analysis.

Although the analysis was performed on a single tubulin-enriched biological sample without replicate LC–MS/MS runs, each reported PTM was supported by multiple peptide-spectrum matches (PSMs) within the acquired dataset. To reflect the total spectral evidence supporting each modified site, the PSM values reported in **Table I** correspond to the sum of all peptide variants carrying the same modification, including different oxidation states and missed-cleavage variants. Although PSM counts provide an indication of the frequency with which modified peptides were detected, they should not be interpreted as quantitative measurements of PTM abundance or site occupancy. High-confidence PTM assignments were obtained using stringent filtering criteria, including a minimum of two PSMs per site, high XCorr scores (up to 7.47), and excellent mass accuracy (DeltaM < 5 ppm). To further support the reliability of these identifications, the highest-quality peptide-spectrum match for each PTM, selected based on XCorr score and unambiguous site localization, is provided in **Supplementary Table S1** together with scan numbers, DeltaScores, and additional identification metrics.

**Table I.**
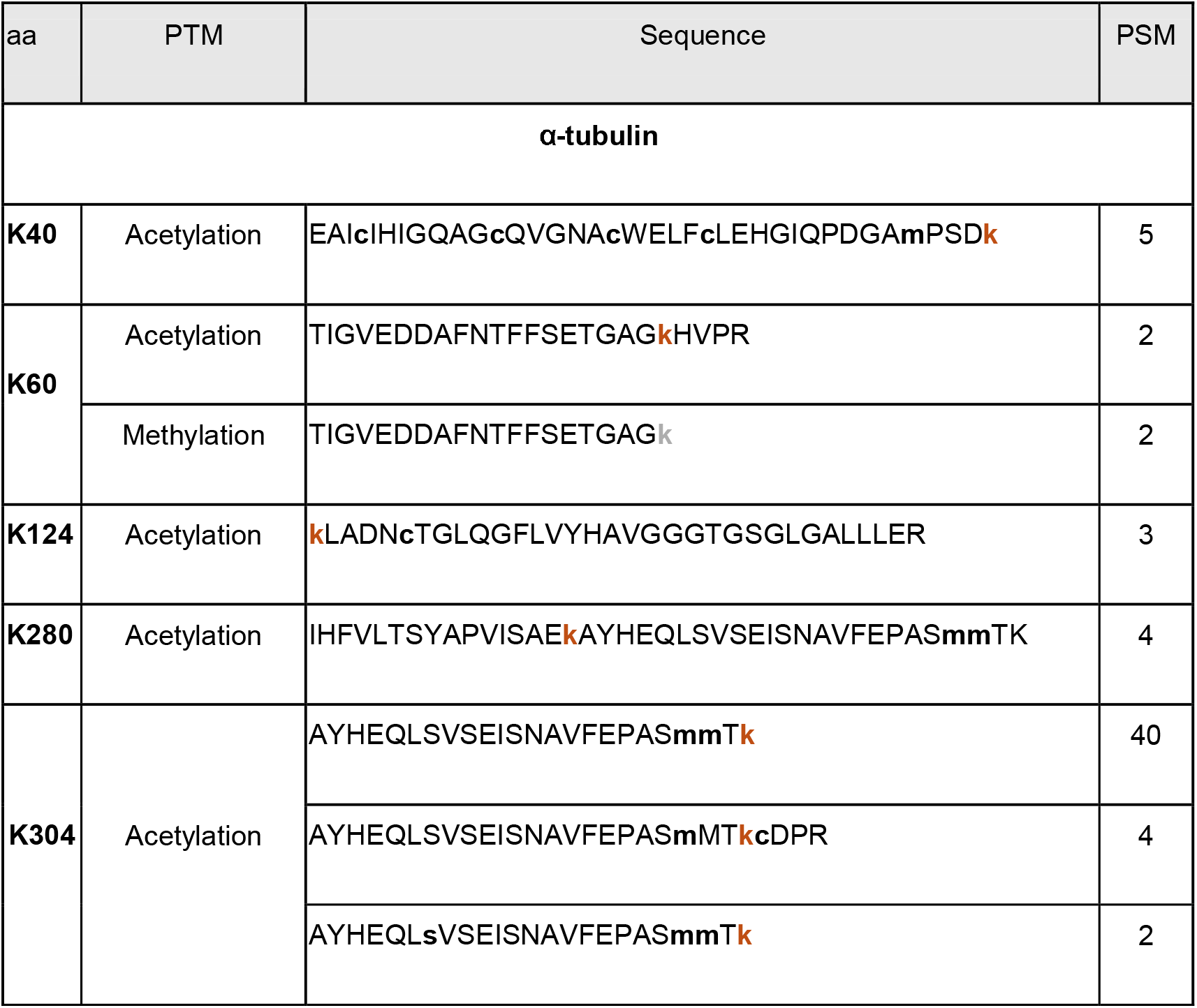
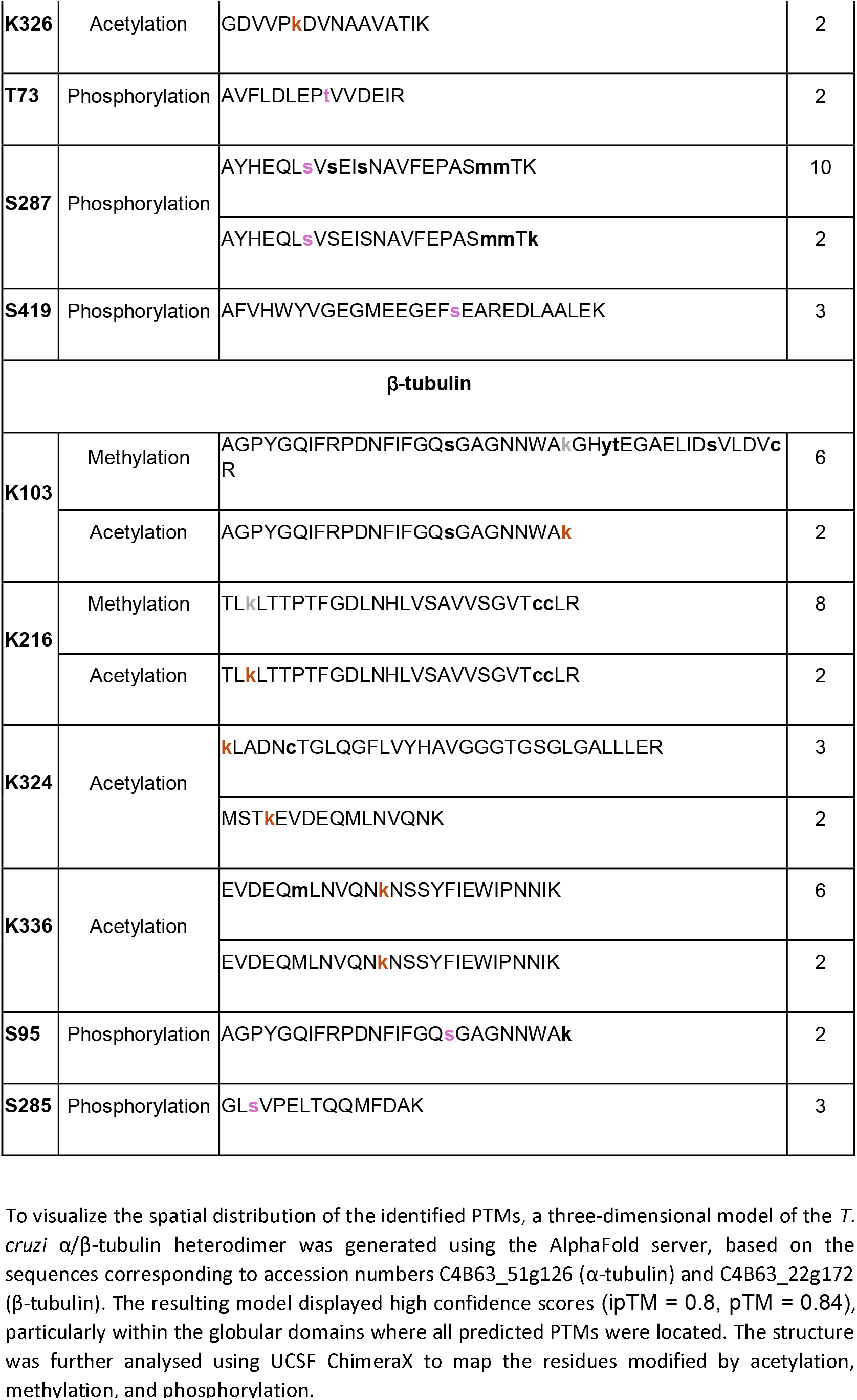
PTMs identified in T. cruzi α- and β-tubulin. Each amino acid residue (aa) is associated with one or more modifications, along with the corresponding peptide sequences and their PSM values. In the peptide sequences, modified amino acids are represented in bold lowercase letters. Specifically, the modified amino acid residue being highlighted in each case is shown in color (orange for acetylation, gray for methylation and violet for phosphorylation), while other PTMs present in the sequence are shown in black bold lowercase.

### Structural modelling

A three-dimensional model of the α/β-tubulin heterodimer was generated using AlphaFold Server, the official portal for AlphaFold3 (RRID:SCR_028034) (Abramson et al., 2024) and visualized with UCSF ChimeraX (RRID:SCR_015872) (v1.11.1) (Pettersen et al., 2021) to map PTM distribution.

## RESULTS

### Purification of *in vitro* polymerized tubulin

Tubulin-enriched extracts were obtained using an *in vitro* polymerization approach based on the addition of GTP and Taxol. The overall workflow of the purification procedure is summarized in **Figure 1A**. Briefly, *T. cruzi* epimastigotes were lysed and subjected to ultracentrifugation to separate the soluble fraction containing MTs (S1) from the insoluble fraction (P0). Tubulin polymerization was then induced in the soluble fraction by incubation with Taxol and GTP, followed by a second ultracentrifugation step to isolate polymerized MTs (P-MT).

**Figure 1.**
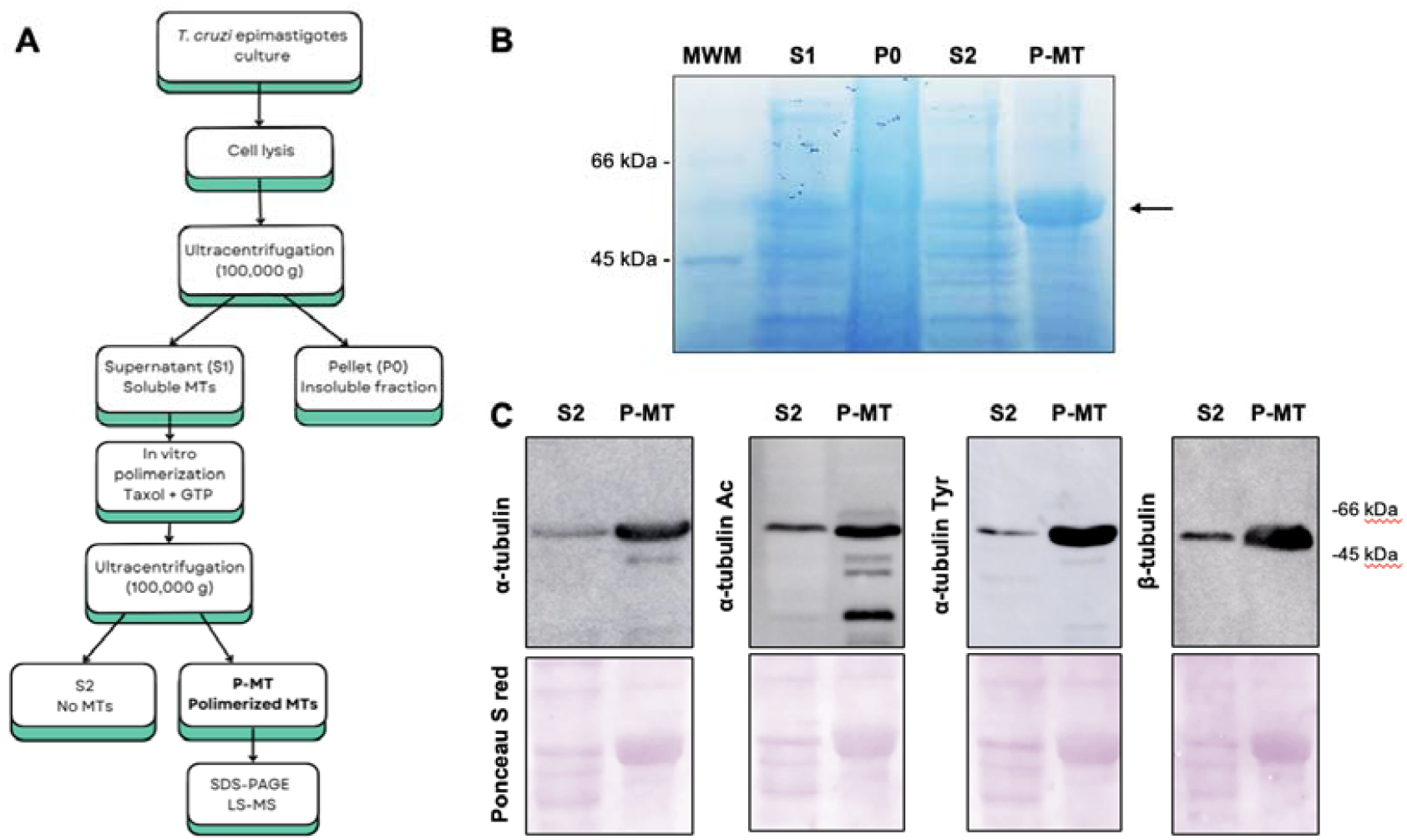
A) Workflow for tubulin enrichment and proteomic analysis. Epimastigotes of *T. cruzi* Dm28c were lysed and the soluble fraction containing MTs (S1) was obtained after ultracentrifugation. Tubulin polymerization was induced *in vitro* by addition of Taxol and GTP, followed by a second ultracentrifugation step to isolate polymerized MTs (P-MT). This fraction was used for SDS-PAGE separation and subsequent LC–MS/MS analysis to identify PTMs in α- and β-tubulin. B) SDS-PAGE analysis of the fractions obtained during the purification procedure. Proteins were separated on 12% polyacrylamide gels and stained with Coomassie Blue. The tubulin band is enriched in the polymerized microtubule fraction (P-MT) compared with the other fractions (arrow). C) Western blot analysis confirming tubulin enrichment in the P-MT fraction. Membranes were probed with antibodies against α-tubulin, acetylated α-tubulin, tyrosinated α-tubulin, and β-tubulin. A strong signal is observed in the P-MT fraction compared with the S2 fraction lacking polymerized MTs.

SDS-PAGE analysis of the different fractions **(Figure 1B**), with protein molecular weight estimated using a low range marker (PB-L Productos Bio-Lógicos), showed a clear enrichment of tubulin in the pellet containing polymerized MTs (P-MT) compared with the other fractions obtained during the purification procedure. Consistently, Western blot analysis (**Figure 1C**) revealed a markedly higher tubulin signal in the P-MT fraction compared with the supernatant lacking polymerized MTs (S2). This enrichment was observed using antibodies against α-tubulin, acetylated α-tubulin, tyrosinated α-tubulin, and β-tubulin. Based on the efficient enrichment of tubulin in this fraction, the P-MT sample was selected for subsequent proteomic analysis by mass spectrometry.

### Identification of PTMs in α- and β-tubulin globular domains

Mass spectrometry results were processed to identify and quantify peptides and proteins present in the samples (Zhu et al., 2010). The datasets obtained using Proteome Discoverer allowed the high-confidence identification of PTMs along the sequences of α- and β-tubulin from *T. cruzi* Dm28c. To ensure high confidence in PTMs identification a stringent filtering criteria were applied: a) only modifications detected in peptides with at least two peptide-spectrum matches (PSMs) were considered; b) PTMs were required to have unambiguous amino acid localization and a mass error (DeltaM) lower than 10 ppm.

The search parameters included both static and dynamic modifications and the dynamic ones were limited to a maximum of one identical modification per peptide and a maximum of four dynamic modifications per peptide. The following dynamic PTMs were included: oxidation, acetylation, methylation, phosphorylation, and tyrosination. N-terminal modifications included: acetylation, methionine loss, and the combined modification methionine loss plus acetylation. Carbamidomethylation was set as a static modification due to the alkylation treatment with iodoacetamide.

The residues in which high-confidence PTMs were detected, together with the type of modification, peptide sequences, and corresponding PSM values, are summarized in **Table I**. Detailed spectral information, mass precision, and localization scores for the best representative match of each identified site are provided in **Supplementary Table S1**.

To visualize the spatial distribution of the identified PTMs, a three-dimensional model of the *T. cruzi* α/β-tubulin heterodimer was generated using the AlphaFold server, based on the sequences corresponding to accession numbers C4B63_51g126 (α-tubulin) and C4B63_22g172 (β-tubulin). The resulting model displayed high confidence scores (ipTM = 0.8, pTM = 0.84), particularly within the globular domains where all predicted PTMs were located. The structure was further analysed using UCSF ChimeraX to map the residues modified by acetylation, methylation, and phosphorylation.

PTMs were identified in α-tubulin, including six acetylation sites (K40, K60, K124, K280, K304, and K326), three phosphorylation sites (T73, S287, and S419), and one methylation site (K60). In β-tubulin, eight PTMs were detected, comprising four acetylation sites (K103, K216, K324, and K336), two phosphorylation sites (S95 and S285), and two methylation sites (K103 and K216) (**Figure 2**). Consistent with the proteomic identification of tubulin methylation, immunoblotting with an anti-methyl-lysine antibody revealed a reactive band at approximately 50 kDa, while immunofluorescence analysis showed partial colocalization of the methyl-lysine signal with tubulin-positive structures in *T. cruzi* epimastigotes (**Supplementary Figure S1**). Quantitative colocalization analysis using the Coloc2 plugin in Fiji revealed a strong positive correlation between both fluorescence channels (Pearson’s correlation coefficient = 0.80 ⍰ 0.02, n = 60 parasites), with Costes significance test indicating that the observed colocalization was significantly greater than expected by chance (P = 1.0 in both analyses).

**Figure 2.**
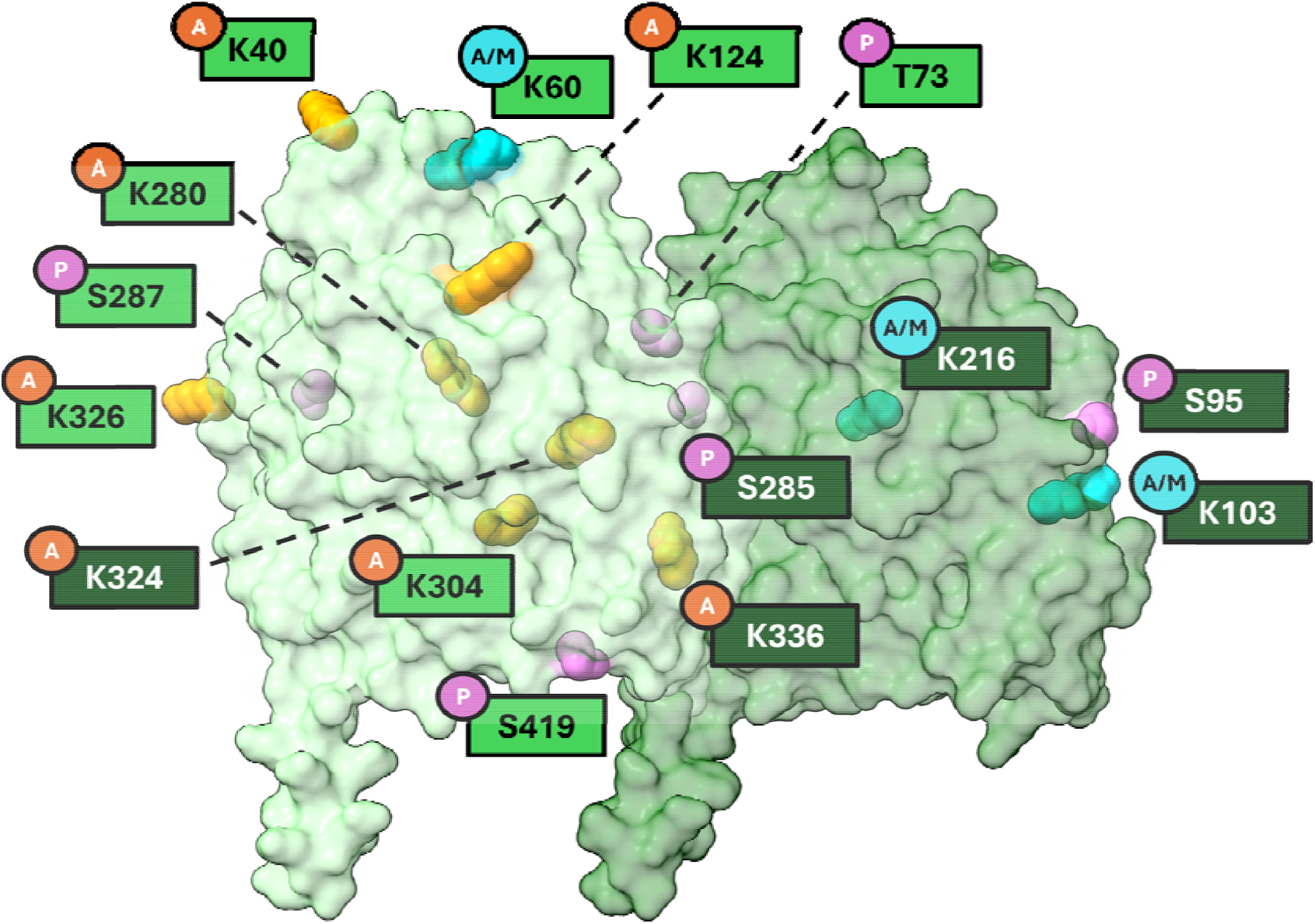
Structural mapping of PTMs on the α/β-tubulin heterodimer of *T. cruzi*. The predicted α/β-tubulin heterodimer is shown based on the AlphaFold model. Both subunits are displayed in a semi-transparent surface representation to allow visualization of modifications: α-tubulin is represented in light green (left) and β-tubulin in dark green (right). Identified PTMs are represented as solid spheres coloured by type: acetylation (orange), phosphorylation (violet). Residues identified with both acetylation and methylation are shown in cyan (αK60, βK103, and βK216). The structural model is intended to illustrate the spatial distribution of PTMs and does not imply experimentally validated functional roles for individual modified residues.

A comparative summary of all tubulin PTMs reported in trypanosomatids, together with the experimental evidence supporting each modification, is provided in **Supplementary Table S2**.

Mapping these PTMs onto the structural model revealed that most PTMs are located on solvent-exposed regions of the tubulin dimer, consistent with potential accessibility to modifying enzymes and interacting proteins, primarily within the globular domains rather than the C-terminal tails. This spatial distribution suggests that these modifications may regulate tubulin function not only through tail-mediated interactions but also by modulating structural regions of the heterodimer that are accessible to modifying enzymes and interacting proteins.

### Polyglutamylation and polyglycylation of the α-tubulin C-terminal tail

Polyglutamylation and polyglycylation were analysed using MaxQuant by performing targeted searches directly on the tubulin sequences. The β-tubulin C-terminal tail could not be confidently analysed under the experimental conditions used, as tryptic digestion generated peptides that were too long for efficient LC–MS/MS characterization.

In contrast, analysis of α-tubulin C-terminal tail enabled the high-confidence detection of polyglutamylation where the search parameters allowed the addition of one, two, or three glutamate residues per modification site. The results are summarized in **Table II**, showing that the primary modified residue corresponds to glutamate E445, followed by E443 and E446. These findings suggest that α-tubulin in *T. cruzi* may exist in multiple isoforms differing in the position or extent of polyglutamylation within the C-terminal tail.

**Table II:** Polyglutamylations identified in *T. cruzi* α-tubulin C-terminal tail. Each amino acid residue (aa) i s associated with one or more modifications, along with the corresponding peptide sequences and their PSM values. In the sequences, modified amino acids are in bold lowercase. Specifically, the residue of interest for polyglutamylation is shown in red, while other PTMs (including additional polyglutamylations) are shown in black bold lowercase.

| aa | Sequence | MS/MS count |
| --- | --- | --- |
| **α-tubulin (C-terminal tail)** |  |  |
| **E443** | SADMe**(1)**GEEDVEEY | 9 |
|  | SADMe**(1)**Ge**(1)**EDVEEY | 14 |
| **E445** | SADmEGe**(2)**EDVEEY | 2 |
|  | SADMEGe**(1)**EDVEEY | 2 |
|  | SADMe**(1)**Ge**(1)**EDVEEY | 14 |
|  | SADMEGe**(1)**e**(1)**DVEEY | 11 |
|  | SADMEGe**(1)**e**(2)**DVEEY | 3 |
| E446 | SADMEGe(1)e(1)DVEEY | 11 |
|  | SADMEGe(1)e(2)DVEEY | 3 |

In contrast, the results obtained for polyglycylation were not robust enough to be considered conclusive. Most peptide identifications presented low MS/MS counts (MS/MS count = 1) and undefined mass error values so the presence of polyglycylation in *T. cruzi* tubulins could not be confidently established. This observation is consistent with previous reports indicating the absence of polyglycylation in other trypanosomatids (Sinclair & de Graffenried, 2019).

### Proposed *T. cruzi* tubulin PTMs landscape

All the information obtained from the proteomic analysis was integrated to generate a representative model of the tubulin PTM landscape in *T. cruzi* (**Figure 3**). This model integrates the distribution of PTMs across both α- and β-tubulin subunits, including PTMs located in the globular domains as well as those detected in the C-terminal tail.

**Figure 3.**
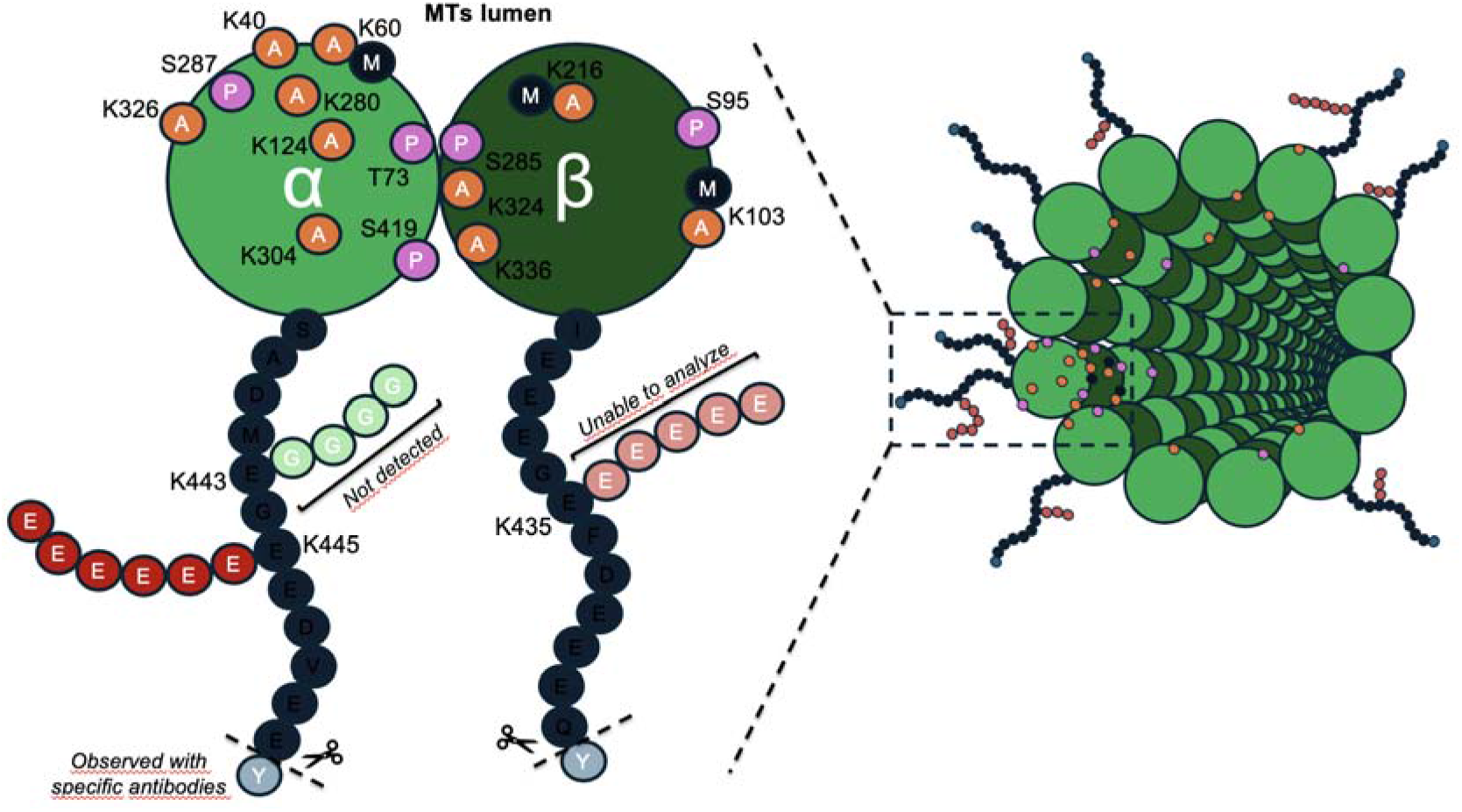
Representative model summarizing detected PTMs. MTs are composed of α- and β-tubulin heterodimers that assemble into protofilaments forming the microtubule lattice. Each tubulin subunit contains a globular domain and a C-terminal tail. The model summarizes the PTMs identified in this study, including acetylation (A, orange), phosphorylation (P, violet), methylation (M, gray), polyglutamylation (E, red), and polyglycylation (G, green). Tyrosination (Y, blue) was included since it was detected using specific antibodies.

The resulting scheme highlights the coexistence of multiple acetylation, phosphorylation, methylation, and polyglutamylation sites, illustrating the potential complexity of the tubulin code operating in *T. cruzi* MTs. Multiple sequence alignment demonstrated that all modified residues identified in α- and β-tubulin are conserved in *T. brucei, Leishmania major* and human tubulins (**Supplementary Figure S2**).

## DISCUSSION

The present study provides the first high-confidence proteomic map of post-translational modifications (PTMs) in α- and β-tubulin from *T. cruzi*. Our analysis identified multiple PTMs, including acetylation, phosphorylation, methylation, and polyglutamylation, distributed across both tubulin subunits. Most of these modification sites had not previously been reported in *T*.

*cruzi*, substantially expanding the known repertoire of tubulin PTMs in this parasite. Sequence alignment of α- and β-tubulins from *T. cruzi, T. brucei, Leishmania major*, and human demonstrated that all modified residues identified in this study are evolutionarily conserved (**Supplementary Figure S2**). Nevertheless, comparison with the available proteomic literature indicates that many of these PTMs have only been experimentally reported in *T. cruzi*, suggesting that species-specific differences arise primarily from PTM regulation rather than divergence of the tubulin primary sequence.

Structural mapping of the identified PTMs revealed distinct topological distributions within the tubulin heterodimer. The acetylated αK40 residue is predicted to face the microtubule lumen, consistent with its established role as a canonical luminal modification associated with microtubule stability in other eukaryotes. In contrast, the polyglutamylated residues identified in the α-tubulin C-terminal tail (E443, E445, and E446) are expected to be exposed on the outer surface of microtubules, where they can regulate interactions with microtubule-associated proteins and motor complexes. Several additional PTMs (αT73, αK280, αS419, βK324, and βS285) are located near the predicted α/β heterodimer interface, raising the possibility that they influence heterodimer stability or conformational dynamics in addition to mediating external protein interactions. These structural assignments are based on AlphaFold modelling and should therefore be considered topological predictions rather than direct evidence of PTM function. Nevertheless, they provide a useful framework for future studies investigating the functional consequences of tubulin PTMs in *T. cruzi*.

The identification of multiple acetylation sites in α- and β-tubulin suggests that acetylation plays a broader regulatory role in the *T. cruzi* cytoskeleton than previously recognized. In trypanosomatids, acetylation of α-tubulin K40 is well established and has been associated with the stability of the subpellicular microtubule array. Our results confirm this conserved modification while identifying five additional acetylation sites in α-tubulin (K60, K124, K280, K304, and K326) that had not previously been reported in *T. cruzi*. Because the present study was designed for PTM identification rather than quantitative analysis, the detection of these sites should not be interpreted as evidence of comparable occupancy or stoichiometry. Among them, K124 and K326 have previously been identified in *Trypanosoma brucei* (Moretti et al., 2018), indicating that some acetylation sites are conserved across trypanosomatids. We also identified four acetylated residues in β-tubulin (K103, K216, K324, and K336). K324 was previously detected in a *T. cruzi* epimastigote acetylome, whereas K216 and K336 were reported in *T. brucei* (Moretti et al., 2018), further supporting partial conservation of tubulin acetylation patterns within this group. Structural mapping predicts that αK60 is located near residues involved in interprotofilament interactions and oriented towards the microtubule lumen. In mammalian tubulin, K60 contributes to contacts with H283 from adjacent protofilaments (Janke & Magiera, 2020). If acetylated in *T. cruzi*, this residue could therefore influence these interactions and alter microtubule mechanical properties, although this hypothesis is based on structural modelling and extrapolation from mammalian studies and remains to be experimentally tested in trypanosomatids. Interestingly, K60 was also identified as a methylation site, raising the possibility that competing PTMs at this residue contribute to the regulation of microtubule stability and mechanical properties (Janke & Magiera, 2020). In *T. cruzi*, acetylation of α-tubulin K40 is catalysed by the acetyltransferase *Tc*ATAT (Alonso et al., 2021), whereas reversible regulation of tubulin acetylation has been associated with members of the HDAC and Sirtuin families (De Oliveira Santos et al., 2019; de Oliveira Santos et al., 2021; Ritagliati et al., 2015). Recent structural studies have shown that tubulin acetyltransferases establish multivalent interactions with the microtubule lattice to access and modify luminal α-tubulin K40 (Luo et al., 2025). In this context, the additional acetylation sites identified here raise the possibility that *Tc*ATAT has a broader substrate specificity than previously recognized or that additional acetyltransferases participate in regulating tubulin acetylation in trypanosomatids. Likewise, although *T. cruzi* encodes only four histone deacetylases (HDACs) and two sirtuins, considerably fewer than mammalian cells (Peralta et al., 2022), the enzymes responsible for tubulin deacetylation remain unknown. These deacetylases may therefore have broader substrate specificities and regulate multiple cellular targets rather than acting exclusively on tubulin.

Comparison with phosphoproteomic datasets from other trypanosomatids revealed both conserved and species-specific phosphorylation sites. The β-tubulin S285 residue appears to be highly conserved, having been identified across different species and life stages, including *Leishmania* promastigotes, bloodstream and procyclic forms of *T. brucei*, and metacyclic forms of *T. cruzi* (Marchini et al., 2011; Morales et al., 2008; Tsigankov et al., 2013; Urbaniak et al., 2013). Likewise, β-tubulin S95, identified in our epimastigote dataset, has also been reported in *T. brucei* (Nett et al., 2009), suggesting a conserved regulatory role within the *Trypanosoma* genus. In contrast, the α-tubulin phosphorylation sites identified in this study (T73, S287, and S419) have not been reported previously in other trypanosomatids. Evidence from *T. cruzi* indicates that tubulin phosphorylation changes during parasite interaction with extracellular matrix components, suggesting a role in environmental sensing and host–parasite interactions (Mattos et al., 2012). In *T. brucei*, phosphorylation of β-tubulin by the cyclin-dependent kinase CRK2 regulates tubulin abundance and restricts excessive microtubule extension, thereby preserving cytoskeletal architecture during the cell cycle (Lee et al., 2023). A similar role has been proposed in *T. cruzi*, where CRK2-mediated tubulin phosphorylation is required to maintain parasite morphology and complete cytokinesis successfully (De Lima et al., 2017). Together, these observations, combined with the phosphorylation sites identified in the present study and the large repertoire of protein kinases encoded in the *T. cruzi* genome, support the view that tubulin phosphorylation is an important regulatory mechanism controlling cytoskeletal dynamics.

Perhaps the most unexpected finding of this study is the identification of methylation sites in both α- and β-tubulin. To our knowledge, tubulin methylation has not previously been reported in trypanosomatids. We identified one methylated residue in α-tubulin (K60) and two in β-tubulin (K103 and K216), all of which were also detected as acetylation sites. This overlap suggests that these lysines may represent regulatory hotspots where different PTMs coexist or compete. In other eukaryotes, tubulin methylation has been implicated in the regulation of microtubule stability and mitotic progression (Cao et al., 2025; Park, Chowdhury, et al., 2016). The best-characterized example is α-tubulin K40, which can be modified either by acetylation through MEC-17/αTAT1 or by methylation through SETD2 (Kearns et al., 2021; Park, Powell, et al., 2016). Whereas acetylation neutralizes the positive charge of the lysine and is associated with stable microtubules, SETD2-mediated trimethylation preserves the positive charge and i s required for proper spindle formation and cytokinesis (Xie et al., 2021). Additional methyltransferases, such as SET8, and demethylases, including KDM4A, further highlight the dynamic regulation of tubulin methylation in mammalian cells (Cao et al., 2025; Chin et al., 2020). Although the functional significance of tubulin methylation in trypanosomatids remains unknown, our findings suggest that similar regulatory mechanisms may operate in the parasite cytoskeleton. The enzymes responsible for these modifications have not yet been identified in *T. cruzi*. However, bioinformatic analyses have revealed candidate methyltransferases in trypanosomatids, and in *T. brucei*, SET19 has been proposed as a functional counterpart of the human methyltransferase SETD2, although its role in tubulin methylation remains to be established (Staneva et al., 2021).

Our targeted analysis of the α-tubulin C-terminal tail identified polyglutamylation at E445 as the predominant modification site, followed by E443 and E446. In eukaryotes, polyglutamylation is a major regulator of microtubule function, modulating interactions with microtubule-associated proteins (MAPs) and motor proteins (Gadadhar et al., 2017; Janke & Magiera, 2020). Recent studies further demonstrated that TTLL-mediated glutamylation contributes to spindle pole integrity and accurate chromosome segregation, with TTLL13 playing a key role in these processes in mammalian cells (Sun et al., 2025). In trypanosomatids, polyglutamylation has been described in *T. brucei* and *Leishmania*, where it is associated with flagellar function and cytokinesis (Casanova et al., 2015; Schneider et al., 1997). In particular, polyglutamylation of α-tubulin E445 is essential during the procyclic stage of *T. brucei*, as deletion of the polyglutamylase TTLL1 reduces modification at this site and results in severe cytokinesis defects, multinucleated cells, and loss of posterior cell integrity (Jentzsch et al., 2020, 2024). The identification of the same modified residue in *T. cruzi* suggests that similar mechanisms may contribute to parasite morphogenesis and cell division. A limitation of the present study is that only the α-tubulin C-terminal tail could be confidently characterized using our trypsin-based bottom-up workflow. The β-tubulin C-terminal peptide generated by tryptic digestion is considerably longer and therefore poorly suited for conventional LC–MS/MS analysis, preventing reliable characterization of tail-associated PTMs. Future studies employing alternative proteases, such as AspN, GluC, or chymotrypsin, could generate shorter peptides more amenable to mass spectrometric analysis. Likewise, middle-down proteomic approaches combined with electron transfer-based fragmentation methods (ETD or EThcD) should substantially improve the identification and localization of polyglutamylation and other PTMs within tubulin C-terminal tails. In contrast, our data did not provide robust evidence for polyglycylation. This result is consistent with previous biochemical and genomic reports indicating that this modification is absent or extremely rare in trypanosomatids, likely due to the evolutionary loss of tubulin glycylase genes within the TTLL family (Nisavic et al., 2025; Schneider et al., 1997; Sinclair & de Graffenried, 2019).

Although tyrosination was not detected by LC–MS/MS, its presence in *T. cruzi* epimastigote extracts was independently supported by Western blot analysis using anti-tyrosinated α-tubulin antibodies (**Figure 1C**). These results are consistent with previous Ultrastructural Expansion Microscopy studies showing the distribution of tyrosinated α-tubulin throughout the epimastigote cytoskeleton (Alonso, 2022). The absence of this PTM from our proteomic dataset likely reflects the well-recognized limitations of trypsin-based bottom-up proteomics for the analysis of tubulin C-terminal tails, as peptides derived from these regions are often difficult to recover because of their length and sequence composition (Janke & Magiera, 2020). A distinctive feature of trypanosomatids is that both α- and β-tubulin are synthesized with a C-terminal tyrosine residue, unlike most other eukaryotes, where this non-canonical residue is restricted to α-tubulin (Sherwin et al., 1987; Sinclair & de Graffenried, 2019). In *T. brucei* and *Leishmania mexicana*, this terminal tyrosine is removed by the tubulin carboxypeptidase Vasohibin (VASH), and a similar processing mechanism has been proposed for other trypanosomatids (Corrales et al., 2025; Thein et al., 2025; van der Laan et al., 2019).

The PTM landscape described here considerably expands the current knowledge of tubulin regulation in *T. cruzi* and provides a valuable resource for future functional studies. The highly specialized and stable microtubule arrays of trypanosomatids, particularly the subpellicular corset, are likely to require multiple regulatory mechanisms to control their organization and function. The diverse PTMs identified in this study provide candidate regulatory sites that may contribute to the specialization of distinct microtubule populations throughout the parasite life cycle. Although the present work provides a comprehensive qualitative map of tubulin PTMs, it does not address their relative abundance, occupancy, or temporal dynamics. Future quantitative and functional studies will therefore be essential to determine how individual and combinatorial PTMs regulate microtubule organization, cytoskeletal dynamics, and parasite biology.

## CONCLUSIONS

This study provides the first comprehensive high-confidence proteomic map of tubulin PTMs in *Trypanosoma cruzi*. Using LC–MS/MS analysis of tubulin-enriched fractions, we identified multiple PTMs, including acetylation, phosphorylation, methylation, and polyglutamylation, in both α- and β-tubulin, substantially expanding the repertoire of tubulin modifications known in this parasite. These findings enabled the proposal of a representative qualitative model of the *T. cruzi* tubulin code and provide a valuable framework for future studies aimed at elucidating the functional significance of individual PTMs and their contribution to microtubule specialization. Future quantitative and targeted proteomic approaches will be required to determine the stoichiometry, occupancy, and temporal dynamics of these modifications, ultimately leading to a more comprehensive understanding of how the tubulin code regulates cytoskeletal organization and parasite biology.

## ACKNOWLEDGMENTS

Mass spectrometric analysis was performed at the Mass Spectrometry Unit of the Institute of Molecular and Cellular Biology of Rosario (UEM-IBR), Argentina. We would like to thank Dr. German Rosano, Lic. Alejo Cantoia and Dr. Enrique Morales for the support with sample preparation and data interpretation of the MS results. Also, we would like to thank Liliana Rojas for her technical assistance.

## DATA AVAILABILITY STATEMENT

We are in the process of making the raw MS data available through the PRIDE database. Accession numbers will be provided upon acceptance. All other data supporting the findings of this study are available from the corresponding author upon reasonable request.

## AUTHOR CONTRIBUTION

Conceived and designed the experiments: VLA, GMP and ES. Performed the experiments: GMP, DB, LB and ADH. Wrote the manuscript: VLA and GMP. Funding acquisition: VLA. All authors contributed to the revisión of the article and approved the submitted version.

## ETHICAL STATEMENT

No human participants or vertebrate animals were involved in this study. *Trypanosoma cruzi* epimastigotes were maintained and manipulated under standard laboratory conditions following the appropriate biosafety requirements. Therefore, ethical approval was not required for this work.

## REFERENCES

Alonso, V. L. (2022). Ultrastructure Expansion Microscopy (U-ExM) in Trypanosoma cruzi: localization of tubulin isoforms and isotypes. Parasitology Research. 10.1007/S00436-022-07619-Z

Alonso, V. L., Carloni, M. E., Gonçalves, C. S., Martinez Peralta, G., Chesta, M. E., Pezza, A., Tavernelli, L. E., Motta, M. C. M., & Serra, E. (2021). Alpha-Tubulin Acetylation in Trypanosoma cruzi: A Dynamic Instability of Microtubules Is Required for Replication and Cell Cycle Progression. Frontiers in Cellular and Infection Microbiology, 11. 10.3389/fcimb.2021.642271

Berná, L., Rodriguez, M., Chiribao, M. L., Parodi-Talice, A., Pita, S., Rijo, G., Alvarez-Valin, F., & Robello, C. (2018). Expanding an expanded genome: long-read sequencing of Trypanosoma cruzi. Microbial Genomics, 4(5). 10.1099/mgen.0.000177

Cao S, Wang S, Xie X, Tan X, Hu X, Shao F, Liu Y, Zhang X, Cheng H, Diao L, Bao L. KDM4A serves as an α-tubulin demethylase regulating microtubule polymerization and cell mitosis. Sci Adv. 2025 Oct 31;11(44):eadv6637. doi: 10.1126/sciadv.adv6637. Epub 2025 Oct 31. PMID: 41171906; PMCID: PMC12577689.

Casanova, M., de Monbrison, F., van Dijk, J., Janke, C., Pagès, M., & Bastien, P. (2015). Characterisation of polyglutamylases in trypanosomatids. International Journal for Parasitology, 45(2–3), 121–132. 10.1016/j.ijpara.2014.09.005

Chin, H. G., Esteve, P. O., Ruse, C., Lee, J., Schaus, S. E., Pradhan, S., & Hansen, U. (2020). The microtubule-associated histone methyltransferase SET8, facilitated by transcription factor LSF, methylates α-tubulin. Journal of Biological Chemistry, 295(14), 4748–4759. 10.1074/jbc.RA119.010951

Corrales RM, Vincent J, Crobu L, Neish R, Nepal B, Espeut J, Pasquier G, Gillard G, Cazevieille C, Mottram JC, Wetzel DM, Sterkers Y, Rogowski K, Lévêque MF. Tubulin detyrosination shapes Leishmania cytoskeletal architecture and virulence. Proc Natl Acad Sci U S A. 2025 Jan 21;122(3):e2415296122. doi: 10.1073/pnas.2415296122. Epub 2025 Jan 14. PMID: 39808657; PMCID: PMC11761321.

De Lima, A. R., Noris-Suárez, K., Bretaña, A., Contreras, V. T., Navarro, M. C., Pérez-Ybarra, L., & Bubis, J. (2017). Growth arrest and morphological changes triggered by emodin on Trypanosoma cruzi epimastigotes cultivated in axenic medium. Biochimie, 142, 31–40. 10.1016/j.biochi.2017.08.005

De Oliveira Santos, J., Zuma, A. A., De Luna Vitorino, F. N., Da Cunha, J. P. C., De Souza, W., & Motta, M. C. M. (2019). Trichostatin A induces Trypanosoma cruzi histone and tubulin acetylation: Effects on cell division and microtubule cytoskeleton remodelling. Parasitology, 146(4), 543–552. 10.1017/S0031182018001828

de Oliveira Santos, J., Zuma, A. A., de Souza, W., & Motta, M. C. M. (2021). Tubastatin A, a deacetylase inhibitor, as a tool to study the division, cell cycle and microtubule cytoskeleton of trypanosomatids. European Journal of Protistology, 80, 125821. 10.1016/J.EJOP.2021.125821

De Souza, W. (2009). Structural organization of Trypanosoma cruzi. Memórias Do Instituto Oswaldo Cruz, 104 Suppl(May), 89–100. http://www.ncbi.nlm.nih.gov/pubmed/19753463

Gadadhar, S., Bodakuntla, S., Natarajan, K., & Janke, C. (2017). The tubulin code at a glance. Journal of Cell Science, 130(8), 1347–1353. 10.1242/jcs.199471

Gull, K. (1999). The cytoskeleton of trypanosomatid parasites. Annual Review of Microbiology, 53, 629–655.

Janke, C., & Magiera, M. M. (2020). The tubulin code and its role in controlling microtubule properties and functions. In Nature Reviews Molecular Cell Biology (Vol. 21, Number 6, pp. 307–326). Nature Research. 10.1038/s41580-020-0214-3

Jentzsch, J., Sabri, A., Speckner, K., Lallinger-Kube, G., Weiss, M., & Ersfeld, K. (2020). Microtubule polyglutamylation is important for regulating cytoskeletal architecture and motility in Trypanosoma brucei. Journal of Cell Science, 133(18). 10.1242/JCS.248047/266504/AM/MICROTUBULE-POLYGLUTAMYLATION-IS-IMPORTANT-FOR

Jentzsch, J., Wunderlich, H., Thein, M., Bechthold, J., Brehm, L., Krauss, S. W., Weiss, M., & Ersfeld, K. (2024). Microtubule polyglutamylation is an essential regulator of cytoskeletal integrity in Trypanosoma brucei. Journal of Cell Science, 137(3). 10.1242/jcs.261740

Kearns, S., Mason, F. M., Rathmell, W. K., Park, I. Y., Walker, C., Verhey, K. J., & Cianfrocco, M. A. (2021). Molecular determinants for α-tubulin methylation by SETD2. Journal of Biological Chemistry, 297(1). 10.1016/j.jbc.2021.100898

Langousis, G., & Hill, K. L. (2014). Motility and more: The flagellum of Trypanosoma brucei. In Nature Reviews Microbiology (Vol. 12, Number 7, pp. 505–518). Nature Publishing Group. 10.1038/nrmicro3274

Lee, K. J., Zhou, Q., & Li, Z. (2023). CRK2 controls cytoskeleton morphogenesis in Trypanosoma brucei by phosphorylating β-tubulin to regulate microtubule dynamics. PLoS Pathogens, 19(3). 10.1371/journal.ppat.1011270

Li, L., & Yang, X. J. (2015). Tubulin acetylation: Responsible enzymes, biological functions and human diseases. In Cellular and Molecular Life Sciences (Vol. 72, Number 22, pp. 4237–4255). Birkhauser Verlag AG. 10.1007/s00018-015-2000-5

Link, A. J., & LaBaer, J. (2009). In-gel trypsin digest of gel-fractionated proteins. Cold Spring Harbor Protocols, 2009(2). 10.1101/pdb.prot5110

Lu, Y. M. (2025). Reinterpreting the effects of α-tubulin K40 acetylation on microtubule stability and cellular functions. Journal of Cell Science, 138(14). 10.1242/jcs.263431

Luo, J., Lam, W.H., Yu, D. et al. Tubulin acetyltransferases access and modify the microtubule luminal K40 residue through anchors in taxane-binding pockets. Nat Struct Mol Biol 32, 358–368 (2025). 10.1038/s41594-024-01406-3

Macrae, T. H., & Gull, K. (1990). Purification and assembly in vitro of tubulin from Trypanosoma brucei brucei. In Biochem. J (Vol. 265).

Marchini, F. K., de Godoy, L. M. F., Rampazzo, R. C. P., Pavoni, D. P., Probst, C. M., Gnad, F., Mann, M., & Krieger, M. A. (2011). Profiling the Trypanosoma cruzi Phosphoproteome. PLOS ONE, 6(9), e25381. 10.1371/journal.pone.0025381

Mattos, E. C., Schumacher, R. I., Colli, W., & Alves, M. J. M. (2012). Adhesion of Trypanosoma cruzi trypomastigotes to fibronectin or laminin modifies tubulin and paraflagellar rod protein phosphorylation. PloS One, 7(10). 10.1371/journal.pone.0046767

Morales, M. A., Watanabe, R., Laurent, C., Lenormand, P., Rousselle, J. C., Namane, A., & Späth, G. F. (2008). Phosphoproteomic analysis of Leishmania donovani pro- and amastigote stages. Proteomics, 8(2), 350–363. 10.1002/pmic.200700697

Moretti, N. S., Cestari, I., Anupama, A., Stuart, K., & Schenkman, S. (2018). Comparative Proteomic Analysis of Lysine Acetylation in Trypanosomes. Journal of Proteome Research, 17(1), 374–385. 10.1021/acs.jproteome.7b00603

Nett, I. R. E., Martin, D. M. A., Miranda-Saavedra, D., Lamont, D., Barber, J. D., Mehlert, A., & Ferguson, M. A. J. (2009). The Phosphoproteome of Bloodstream Form Trypanosoma brucei, Causative Agent of African Sleeping Sickness. Molecular & Cellular Proteomics, 8(7), 1527–1538. 10.1074/mcp.M800556-MCP200

Nisavic M, Chaze T, Bonnefoy S, Janke C, Bastin P, Matondo M, Chamot-Rooke J. Mass Spectrometry Reveals Novel Features of Tubulin Polyglutamylation in the Flagellum of Trypanosoma brucei. J Proteome Res. 2025 Aug 1;24(8):3979–3989. doi: 10.1021/acs.jproteome.5c00107. Epub 2025 Jul 9. PMID: 40634267.

Orsburn, B. C. (2021). Proteome Discoverer-A Community Enhanced Data Processing Suite for Protein Informatics. Proteomes, 9(1). 10.3390/proteomes9010015

Park, I. Y., Chowdhury, P., Tripathi, D. N., Powell, R. T., Dere, R., Terzo, E. A., Rathmell, W. K., & Walker, C. L. (2016). Methylated α-tubulin antibodies recognize a new microtubule modification on mitotic microtubules. MAbs, 8(8), 1590–1597. 10.1080/19420862.2016.1228505

Park, I. Y., Powell, R. T., Tripathi, D. N., Dere, R., Ho, T. H., Blasius, T. L., Chiang, Y. C., Davis, I. J., Fahey, C. C., Hacker, K. E., Verhey, K. J., Bedford, M. T., Jonasch, E., Rathmell, W. K., & Walker, C. L. (2016). Dual Chromatin and Cytoskeletal Remodeling by SETD2. Cell, 166(4), 950–962. 10.1016/j.cell.2016.07.005

Peralta, G. M., Serra, E., & Alonso, V. L. (2022). Update on the Biological Relevance of Lysine Acetylation as a Novel Drug Target in Trypanosomatids. Current Medicinal Chemistry, 29(20), 3638–3659. 10.2174/0929867328666211126145721

Perdiz, D., Mackeh, R., Poüs, C., & Baillet, A. (2011). The ins and outs of tubulin acetylation: More than just a post-translational modification? In Cellular Signalling (Vol. 23, Number 5, pp. 763–771). 10.1016/j.cellsig.2010.10.014

Ritagliati, C., Alonso, V. L., Manarin, R., Cribb, P., & Serra, E. C. (2015). Overexpression of Cytoplasmic TcSIR2RP1 and Mitochondrial TcSIR2RP3 Impacts on Trypanosoma cruzi Growth and Cell Invasion. PLoS Neglected Tropical Diseases, 9(4). 10.1371/journal.pntd.0003725

Schneider, A., Plessmann, U., & Weber, K. (1997). Subpellicular and flagellar microtubules of Trypanosoma brucei are extensively glutamylated. Journal of Cell Science, 110(4), 431–437. 10.1242/jcs.110.4.431

Sherwin, T., Schneider, A., Sasse, R., Seebeck, T., & Gull, K. (1987). Distinct localization and cell cycle dependence of COOH terminally tyrosinated alpha-tubulin in the microtubules of Trypanosoma brucei brucei. J. Cell Biol., 104(March), 439–446. http://www.ncbi.nlm.nih.gov/entrez/query.fcgi?cmd=Retrieve&db=PubMed&dopt=Citation&list_uids=3546334

Sinclair, A. N., & de Graffenried, C. L. (2019). More than Microtubules: The Structure and Function of the Subpellicular Array in Trypanosomatids. In Trends in Parasitology (Vol. 35, Number 10, pp. 760–777). Elsevier Ltd. 10.1016/j.pt.2019.07.008

Staneva, D. P., Carloni, R., Auchynnikava, T., Tong, P., Rappsilber, J., Arockia Jeyaprakash, A., Matthews, K. R., & Allshire, R. C. (2021). A systematic analysis of Trypanosoma brucei chromatin factors identifies novel protein interaction networks associated with sites of transcription initiation and termination. Genome Research, 31(11), 2138–2154. 10.1101/GR.275368.121/-/DC1

Sun, S., Wang, Z., Xu, Z. et al. CFAP100 couples microtubule glutamylation to spindle pole integrity in keratinocytes to promote epidermal development. Nat Commun 16, 5591 (2025). 10.1038/s41467-025-60677-6

Thein M, Wunderlich H, Brehm L, Wagner S, Weiss M, Ersfeld K. Effects of microtubule (de)tyrosination on the morphology and motility of Trypanosoma brucei and cross-talk with polyglutamylation. Biol Open. 2025 Dec 15;14(12):bio062270. doi: 10.1242/bio.062270. Epub 2025 Dec 16. PMID: 41400423; PMCID: PMC12714142.

Tsigankov, P., Gherardini, P. F., Helmer-Citterich, M., Späth, G. F., & Zilberstein, D. (2013). Phosphoproteomic analysis of differentiating Leishmania parasites reveals a unique stage-specific phosphorylation motif. Journal of Proteome Research, 12(7), 3405–3412. 10.1021/pr4002492

Urbaniak, M. D., Martin, D. M. A., & Ferguson, M. A. J. (2013). Global quantitative SILAC phosphoproteomics reveals differential phosphorylation is widespread between the procyclic and bloodstream form lifecycle stages of Trypanosoma brucei. Journal of Proteome Research, 12(5), 2233–2244. 10.1021/pr400086y

van der Laan, S., Lévêque, M. F., Marcellin, G., Vezenkov, L., Lannay, Y., Dubra, G., Bompard, G., Ovejero, S., Urbach, S., Burgess, A., Amblard, M., Sterkers, Y., Bastien, P., & Rogowski, K. (2019). Evolutionary Divergence of Enzymatic Mechanisms for Tubulin Detyrosination. Cell Reports, 29(12), 4159–4171.e6. 10.1016/J.CELREP.2019.11.074

Vidal, J. C., & Souza, W. de. (2017). Morphological and Functional Aspects of Cytoskeleton of Trypanosomatids. In Cytoskeleton - Structure, Dynamics, Function and Disease. InTech. 10.5772/66859

Xie, X., Wang, S., Li, M., Diao, L., Pan, X., Chen, J., Zou, W., Zhang, X., Feng, W., & Bao, L. (2021). α-TubK40me3 is required for neuronal polarization and migration by promoting microtubule formation. Nature Communications, 12(1). 10.1038/s41467-021-24376-2

Zhu, W., Smith, J. W., & Huang, C. M. (2010). Mass spectrometry-based label-free quantitative proteomics. Journal of Biomedicine & Biotechnology, 2010. 10.1155/2010/840518

Zingales, B., Miles, M. A., Campbell, D. A., Tibayrenc, M., Macedo, A. M., Teixeira, M. M. G., Schijman, A. G., Llewellyn, M. S., Lages-Silva, E., Machado, C. R., Andrade, S. G., & Sturm, N. R. (2012). The revised Trypanosoma cruzi subspecific nomenclature: Rationale, epidemiological relevance and research applications. In Infection, Genetics and Evolution (Vol. 12, Number 2, pp. 240–253). 10.1016/j.meegid.2011.12.009

Zuma, A. A., dos Santos Barrias, E., & de Souza, W. (2021). Basic Biology of Trypanosoma cruzi. Current Pharmaceutical Design, 27(14), 1671–1732. 10.2174/1381612826999201203213527

