## Supplementary Figure 1,2 and Table 1 for "Proteomic mapping of novel tubulin post-translational modifications in *Trypanosoma cruzi* cytoskeleton"

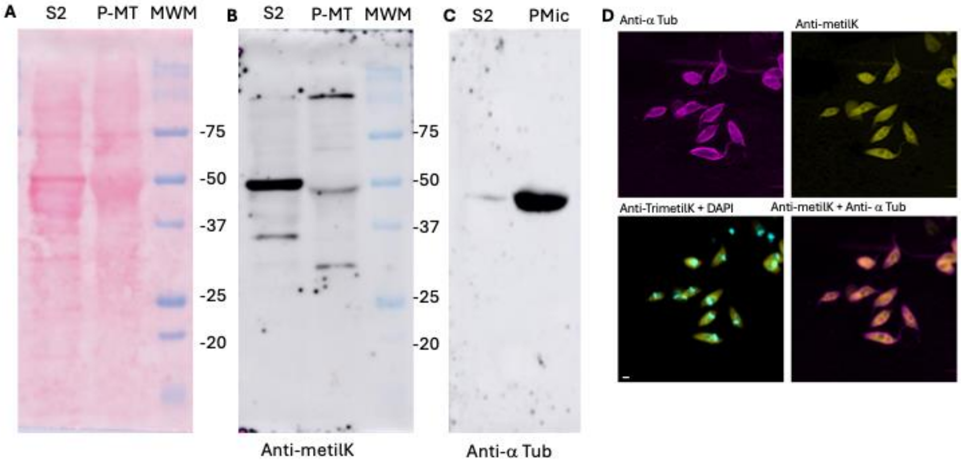


**Supplementary Figure S1: Experimental validation of tubulin methylation in *Trypanosoma cruzi* epimastigotes.** Tubulin-enriched polymerized microtubule fraction (PMic) and the corresponding soluble fraction (S2) were analyzed by Western blot and indirect immunofluorescence. (A) Ponceau S staining of nitrocellulose membranes showing total protein loading. (B) Western blot using an anti-trimethyl-lysine antibody. A prominent immunoreactive band at the expected molecular weight of tubulin was detected in the PMic fraction, together with additional methylated proteins. (C) Western blot using an anti-α-tubulin antibody confirming the enrichment of tubulin in the PMic fraction. MWM: Molecular Wight Marker. (D) Representative indirect immunofluorescence images of epimastigotes stained with anti-α-tubulin (magenta), anti-methyl-lysine (yellow), DAPI (cyan), and the corresponding merged images. Bar = 10 µm.

**A** CLUSTAL multiple sequence alignment by MUSCLE (3.8) Alpha-tubulin

```
Human           MRECISIHVGQAGVQIGNACWELYCLEHGIQPDGQMPSDKTIGGGDDSFNTFFSETGAGK
T.cruzi         MREAICIHIGQAGCQVGNACWELFCLEHGIQPDGAMPSDKTIGVEDDAFNTFFSETGAGK
T.brucei        MREAICIHIGQAGCQVGNACWELFCLEHGIQPDGAMPSDKTIGVEDDAFNTFFSETGAGK
L.major         MREAICIHIGQAGCQVGNACWELFCLEHGIQPDGSMPSDKCIGVEDDAFNTFFSETGAGK
                ***.*.**:**** *:*******:********** ***** **  **:************

Human           HVPRAVFVDLEPTVIDEVRTGTYRQLFHPEQLITGKEDAANNYARGHYTIGKEIIDLVLD
T.cruzi         HVPRAVFLDLEPTVVDEIRTGTYRQLFHPEQLISGKEDAANNYARGHYTIGKEIVDLCLD
T.brucei        HVPRAVFLDLEPTVVDEVRTGTYRQLFHPEQLISGKEDAANNYARGHYTIGKEIVDLCLD
L.major         HVPRCIFLDLEPTVVDEVRTGTYRQLFNPEQLVSGKEDAANNYARGHYTIGKEIVDLALD
                ****.:*:******:**:*********:****::********************:** **

Human           RIRKLADQCTGLQGFLVFHSFGGGTGSGFTSLLMERLSVDYGKKSKLEFSIYPAPQVSTA
T.cruzi         RIRKLADNCTGLQGFLVYHAVGGGTGSGLGALLLERLSVDYGKKSKLGYTVYPSPQVSTA
T.brucei        RIRKLADNCTGLQGFLVYHAVGGGTGSGLGALLLERLSVDYGKKSKLGYTVYPSPQVSTA
L.major         RIRKLADNCTGLQGFMVFHAVGGGTGSGLGALLLERLSVDYGKKSKLGYTVYPSPQVSTA
                *******:*******:*:*:.*******: :**:************* :::**:******

Human           VVEPYNSILTTHTTLEHSDCAFMVDNEAIYDICRRNLDIERPTYTNLNRLISQIVSSITA
T.cruzi         VVEPYNSVLSTHSLLEHTDVAAMLDNEAIYDLTRRNLDIERPTYTNLNRLIGQVVSALTA
T.brucei        VVEPYNSVLSTHSLLEHTDVAAMLDNEAIYDLTRRNLDIERPTYTNLNRLIGQVVSSLTA
L.major         VVEPYNCVLSTHSLLEHTDVATMLDNEAIYDLTRRSLDIERPSYTNVNRLIGQVVSSLTA
                ******.:*:**: ***:* * *:*******: **.******:***:****.*:**::**

Human           SLRFDGALNVDLTEFQTNLVPYPRIHFPLATYAPVISAEKAYHEQLSVAEITNACFEPAN
T.cruzi         SLRFDGALNVDLTEFQTNLVPYPRIHFVLTSYAPVISAEKAYHEQLSVSEISNAVFEPAS
T.brucei        SLRFDGALNVDLTEFQTNLVPYPRIHFVLTSYAPVISAEKAYHEQLSVSEISNAVFEPAS
L.major         SLRFDGALNVDLTEFQTNLVPYPRIHFVLTSYAPVVSAEKAYHEQLSVADITNSVFEPAG
```

```
                     ***************************  *::****:************::*:*: ****.

Human                QMVKCDPRHGKYMACCLLYRGDVVPKDVNAAIATIKTKRSIQFVDWCPTGFKVGINYQPP
T.cruzi              MMTKCDPRHGKYMACCLMYRGDVVPKDVNAAVATIKTKRTIQFVDWSPTGFKCGINYQPP
T.brucei             MMTKCDPRHGKYMACCLMYRGDVVPKDVNAAVATIKTKRTIQFVDWSPTGFKCGINYQPP
L.major              MLTKCDPRHGKYMSCCLMYRGDVVPKDVNAAIATIKTKRTIQFVDWCPTGFKCGINYQPP
                      :.**********:***:*************:*******:******.***** *******

Human                TVVPGGDLAKVQRAVCMLSNTTAIAEAWARLDHKFDLMYAKRAFVHWYVGEGMEEGEFSE
T.cruzi              TVVPGGDLAKVQRAVCMIANSTAIAEVFARIDHKFDLMYSKRAFVHWYVGEGMEEGEFSE
T.brucei             TVVPGGDLAKVQRAVCMIANSTAIAEVFARIDHKFDLMYSKRAFVHWYVGEGMEEGEFSE
L.major              TVVPGGDLAKVQRAVCMIANSTAIAEVFARIDHKFDLMYSKRAFVHWYVGEGMEEGEFSE
                     *****************::*:*****.:**:********:********************

Human                AREDMAALEKDYEEVGVDSVEGEGEEEGEEY
T.cruzi              AREDLAALEKDYEEVGAESADMEGEEDVEEY
T.brucei             AREDLAALEKDYEEVGAESADMDGEEDVEEY
L.major              AREDLAALEKDYEEVGAESADDMGEEDVEEY
                     ****:***********.:*.:  ***: ***
```

**B** CLUSTAL multiple sequence alignment by MUSCLE (3.8) Beta-tubulin

```
Human                MREIVHIQIGQCGNQIGAKFWEMIGEEHGIDLAGSDRGASALQLERISVYYNEAYGRKYV
L.major              MREIVSCQAGQCGNQIGSKFWEVISDEHGVDPTGTYQGDSDLQLERINVYFDESTGGRYV
T.cruzi              MREIVCVQAGQCGNQIGSKFWEVISDEHGVDPTGTYQGDSDLQLERINVYFDEATGGRYV
T.brucei             MREIVCVQAGQCGNQIGSKFWEVISDEHGVDPTGTYQGDSDLQLERINVYFDEATGGRYV
                     *****  * ********:****:*.:***:* :*: .* * ******.**::*: * .**

Human                PRAVLVDLEPGTMDSIRSSKLGALFQPDSFVHGNSGAGNNWAKGHYTEGAELIENVLEVV
L.major              PRAVLMDLEPGTMDSVRAGPYGQLFRPDNFIFGQSGAGNNWAKGHYTEGAELIDSVLDVC
T.cruzi              PRAVLIDLEPGTMDSVRAGPYGQIFRPDNFIFGQSGAGNNWAKGHYTEGAELIDSVLDVC
T.brucei             PRSVLIDLEPGTMDSVRAGPYGQIFRPDNFIFGQSGAGNNWAKGHYTEGAELIDSVLDVC
                     **:**:*********:*:.  * :*.**.*:.*:*******************:.**:* 

Human                RHESESCDCLQGFQIVHSLGGGTGSGMGTLLMNKIREEYPDRIMNSFSVMPSPKVSDTVV
L.major              RKEAESCDCLQGFQLSHSLGGGTGSGMGTLLISKLREEYPDRIMMTFSVIPSPRVSDTVV
T.cruzi              RKEAESCDCLQGFQICHSLGGGTGSGMGTLLISKLREEYPDRIMMTFSIIPSPKVSDTVV
T.brucei             CKEAESCDCLQGFQICHSLGGGTGSGMGTLLISKLREQYPDRIMMTFSIIPSPKVSDTVV
                      :*:**********: ***************:.*:**:****** :**::***.******

Human                EPYNAVLSIHQLIENADACFCIDNEALYDICFRTLKLTTPTYGDLNHLVSLTMSGITTSL
L.major              EPYNTTLSVHQLVENSDESMCIDNEALYDICFRTLKLTTPTFGDLNHLVAAVMSGVTCCL
T.cruzi              EPYNTTLSVHQLVENSDESMCIDNEALYDICFRTLKLTTPTFGDLNHLVSAVVSGVTCCL
T.brucei             EPYNTTLSVHQLVENSDESMCIDNEALYDICFRTLKLTTPTFGDLNHLVSAVVSGVTCCL
                     ****:.**:***:**:* .:*********************:*******: .:**:* .*

Human                RFPGQLNADLRKLAVNMVPFPRLHFFMPGFAPLTAQGSQQYRALSVAELTQQMFDARNTM
L.major              RFPGQLNSDLRKLAVNLVPFPRLHFFMMGFAPLTSRGSQQYRGLSVAELTQQMFDAKNMM
T.cruzi              RFPGQLNSDLRKLAVNLVPFPRLHFFMMGFAPLTSRGSQQYRGLSVPELTQQMFDAKNMM
T.brucei             RFPGQLNSDLRKLAVNLVPFPRLHFFMMGFAPLTSRGSQQYRGLSVPELTQQMFDAKNMM
                     *******:********:********** ******:.******.***.*********.* *

Human                AACDLRRGRYLTVACIFRGKMSTKEVDQQLLSVQTRNSSCFVEWIPNNVKVAVCDIPPRG
L.major              QAADPRHGRYLTASALFRGRMSTKEVDEQMLNVQNKNSSYFIEWIPNNIKSSICDIPPKG
T.cruzi              QAADPRHGRYLTASALFRGRMSTKEVDEQMLNVQNKNSSYFIEWIPNNIKSSICDIPPKG
T.brucei             QAADPRHGRYLTASALFRGRMSTKEVDEQMLNVQNKNSSYFIEWIPNNIKSSVCDIPPKG
                      *.* *.*****.:.:***.*******:*:*.**..*** *:******:* ::*****.*
```

```
Human         LSMAATFIGNNTAIQEIFNRVSEHFSAMFKRKAFVHWYTSEGMDINEFGEAENNIHDLVS
L.major       LKMSVTFIGNNTCIQEMFRRVGEQFTGMFRRKAFLHWYTGEGMDEMEFTEAESNMNDLVS
T.cruzi       LKMAVTFVGNNTCIQEMFRRVGEQFTAMFRRKAFLHWYTGEGMDEMEFTEAESNMNDLVS
T.brucei      LKMAVTFIGNNTCIQEMFRRVGEQFTLMFRRKAFLHWYTGEGMDEMEFTEAESNMNDLVS
              *.*:.**:****.***:*.**.*:*: **.****:****.****  ** ***.*::****

Human         EYQQFQDAKAVLEEDEEVTEEAEMEPEDKGH
L.major       EYQQYQDAT--------VEEEGEYDEEEEAY
T.cruzi       EYQQYQDAT--------IEEEGEFDEEEQ-Y
T.brucei      EYQQYQDAT--------IEEEGEFDEEEQ-Y
              ****:***.        : **.* : *:: :
```

**Supplementary Figure S2:** Multiple sequence alignment of α-tubulin (A) and β-tubulin (B) from Trypanosoma cruzi Dm28c, Trypanosoma brucei, Leishmania major and Homo sapiens generated using MUSCLE. Residues identified as post-translationally modified in the present study are highlighted in color red. Asterisks indicate identical residues among all four species, whereas colons and periods denote conservative and semi-conservative substitutions, respectively. Sequence accession numbers are: Alfa Tubulin *Homo sapiens* TUBA1A P68363, *Trypanosoma cruzi* Dm28c C4B63_51g126, *Trypanosoma brucei* Tb927.1.2340, *Leishmania major* LMJFC_130008200. Beta Tubulin *Homo sapiens* TUBB1 Q9H4B7, *Trypanosoma cruzi* C4B63_22g172, *Trypanosoma brucei* Tb427.01.2330 and *Leishmania major* LMJFC_210030600.

**Supplementary Table S1:** Spectral parameters of identified tubulin PTMs. This table summarizes the high-confidence proteomic parameters for each modification site identified in *T. cruzi* α- and β-tubulin. To ensure maximum data rigor, a best match representative is reported for each residue, corresponding to the individual MS/MS spectrum with the highest XCorr value. Total PSM values represent all peptide variants identified for each specific residue (including different oxidation states or undetected cleavage variants).Additionally, DeltaScore (site localization probability) and DeltaM (ppm) (mass precision) values are provided to confirm unambiguous PTM localization and the high-resolution nature of the mass spectrometry identification.

| Residue | PTM Type | Representative Scan | Total PSMs | Best XCorr | DeltaScore | DeltaM (ppm) |
|---|---|---|---|---|---|---|
| **α-tubulin** | | | | | | |
| K40 | Acetylation | 22336 | 5 | 5.18 | 0.6892 | 4.30 |
| K60 | Acetylation | 23316 | 2 | 6.63 | 0.1041 | -1.88 |
| K60 | Methylation | 27396 | 2 | 4.62 | 0.7511 | -0.40 |
| T73 | Phosphorylation | 30171 | 2 | 5.45 | 0.6404 | 0.47 |
| K124 | Acetylation | 29919 | 3 | 7.24 | 0.8508 | 0.40 |
| K280 | Acetylation | 27427 | 4 | 3.95 | 0.0076 | 2.46 |
| S287 | Phosphorylation | 26989 | 12 | 7.47 | 0.1058 | -0.97 |
| K304 | Acetylation | 20082 | 46 | 4.75 | 0.0189 | 4.35 |
| K326 | Acetylation | 17644 | 2 | 2.97 | 0.5825 | -1.60 |
| S419 | Phosphorylation | 28067 | 3 | 6.48 | 0.5633 | 0.43 |
| **β-tubulin** | | | | | | |
| S95 | Phosphorylation | 23406 | 2 | 5.09 | 0.6660 | -4.22 |
| K103 | Acetylation | 23406 | 2 | 5.09 | 0.6660 | -4.22 |
| K103 | Methylation | 25421 | 6 | 2.80 | 0.0786 | 1.85 |
| K216 | Acetylation | 35090 | 2 | 4.50 | 0.7533 | 1.76 |
| K216 | Methylation | 33561 | 8 | 7.36 | 0.8424 | -3.62 |

| | | | | | | |
|---|---|---|---|---|---|---|
| S285 | Phosphorylation | 27493 | 3 | 4.02 | 0.5000 | -0.78 |
| K324 | Acetylation | 14592 | 5 | 4.04 | 0.7079 | -1.43 |
| K336 | Acetylation | 27399 | 8 | 4.66 | 0.4914 | -3.00 |

**Supplementary Table S2:** Comparative distribution of tubulin PTMs in trypanosomatids. The table summarizes the PTMs identified in α- and β-tubulin in the present study and compares them with those previously reported in *Trypanosoma cruzi*, *Trypanosoma brucei* and *Leishmania* spp. For each modified residue, the type of experimental evidence is indicated as MS (mass spectrometry), Ab (antibody-based detection), Gen (genetic or functional evidence), or NR (not reported). PTMs identified in the present study are indicated as MS (this study). Corresponding references are provided for all previously reported modifications.

| PTM type | Residue (T. cruzi) | T. cruzi | T. brucei | Leishmania spp. | References |
|---|---|---|---|---|---|
| Acetylation | αK40 | MS / Ab / Gen | MS / Ab | Ab | Alonso et al., 2021; Moretti et al., 2018 |
| Acetylation | αK60 | MS (this study) | NR | NR | This study |
| Methylation | αK60 | MS (this study) | NR | NR | This study |
| Phosphorylation | αT73 | MS (this study) | NR | NR | This study |
| Acetylation | αK124 | MS (this study) | MS | NR | Moretti et al., 2018 |
| Acetylation | αK280 | MS (this study) | NR | NR | This study |
| Phosphorylation | αS287 | MS (this study) | NR | NR | This study |
| Acetylation | αK304 | MS (this study) | NR | NR | This study |
| Acetylation | αK326 | MS (this study) | MS | NR | Moretti et al., 2018 |
| Phosphorylation | αS419 | MS (this study) | NR | NR | This study |
| Polyglutamylation | αE443 | MS (this study) | Ab/Gen (C-terminal tail) | Ab (C-terminal tail) | Jentzsch et al., 2020; Schneider et al., 1997 |
| Polyglutamylation | αE445 | MS (this study) | Ab / Gen | Ab | Jentzsch et al., 2020; Schneider et al., 1997 |
| Polyglutamylation | αE446 | MS (this study) | Ab/Gen (C-terminal tail) | Ab (C-terminal tail) | Jentzsch et al., 2020 |
| Phosphorylation | βS95 | MS (this study) | MS | NR | Nett et al., 2009 |
| Acetylation | βK103 | MS (this study) | NR | NR | This study |
| Methylation | βK103 | MS (this study) | NR | NR | This study |
| Acetylation | βK216 | MS (this study) | MS | NR | Moretti et al., 2018 |

| | | | | | |
|---|---|---|---|---|---|
| Methylation | βK216 | MS (this study) | NR | NR | This study |
| Phosphorylation | βS285 | MS | MS | MS | Marchini et al., 2011; Urbaniak et al., 2013; Tsigankov et al., 2013 |
| Acetylation | βK324 | MS | NR | NR | Moretti et al., 2018 |
| Acetylation | βK336 | MS (this study) | MS | NR | Moretti et al., 2018 |
| Tyrosination | α C-terminal Tyr | Ab | Ab / MS | Ab | Schneider et al., 1997; Alonso, 2022 |
| Tyrosination | β C-terminal Tyr | Ab | Ab / MS | Ab | Schneider et al., 1997; Alonso, 2022 |
